# Effect of environment on the evolutionary trajectories and growth characteristics of antibiotic-resistant *Escherichia coli* mutants

**DOI:** 10.1101/262691

**Authors:** Alasdair T. M. Hubbard, Nazila V. Jafari, Nicholas Feasey, Jennifer L. Rohn, Adam P. Roberts

## Abstract

In the face of an accelerating global antimicrobial resistance crisis, the determination of bacterial fitness following acquisition of resistance is an expanding area of research, and increased understanding of this process will be crucial to translate *in vitro* fitness data to successful therapies. Given that crucial clinical treatment situations are guided by *in vitro* diagnostic testing in an artificial environment far removed from human physiological niches, we used *Escherichia coli* and amoxicillin-clavulanic acid (AMC) resistance as a model to understand how such environments could affect the emergence of resistance, associated fitness costs and the predictive value of this data when strains were grown in the more physiologically relevant environments of urine and urothelial organoids. Resistant *E. coli* isolates were selected for following 24-hour exposure to sub-inhibitory concentrations of AMC in either M9, ISO or LB broth, followed by growth on LB agar containing AMC. No resistant colonies emerged following growth in M9, whereas resistant isolates were detected from cultures grown in ISO and LB broth. We observed both within and between media-type variability in the levels of resistance and fitness of the resistant mutants grown in LB. MICs and fitness of these resistant strains in different media (M9, ISO, LB, human urine and urothelial organoids) showed considerable variation. Media can therefore have a direct effect on the isolation of mutants that confer resistance to AMC and these mutants can exhibit unpredictable MIC and fitness profiles under different growth conditions. This study highlights the risks in relying on a single culture protocol to predict the behaviour and treatment response of bacteria *in vivo* and highlights the importance of developing comprehensive experimental designs to ensure effective translation of diagnostic procedures to successful clinical outcomes.

## Introduction

The alarming global rise in antimicrobial resistance (AMR) has led to increased interest in resistance development, persistence and fixation within bacterial populations, with a wide variety of studies published in recent years. These include determining whether the prevalence of resistant bacteria is exacerbated by antimicrobials and their metabolites entering the environment following use in humans [1, 2], understanding the evolution of AMR due to selective pressure from antimicrobials, and establishing how the acquisition of AMR genes affects bacterial fitness [3–5]. Ultimately, such investigations should lead to new strategies to prevent the development and / or persistence of AMR in the future.

From an evolutionary perspective, there is a particular focus on the effect of acquisition of AMR on bacterial fitness with a view to suppress the development and persistence of resistance [4, 6, 7]. When bacteria acquire resistance, either through mutation or horizontal gene transfer, they often incur a fitness cost. Many of these evolutionary studies rely on the *in vitro* development of resistance in the presence of sub-inhibitory concentrations of single, or combinations of, antimicrobials, and investigators employ a range of different media for the generation of resistance [6, 8–10] as well as the subsequent assessment of fitness related to acquired AMR [11, 12]. However, the media currently used to grow bacteria *in vitro* is not necessarily the most suitable substitute for the *in vivo* environment in which the bacteria live. A recent study found that mutations that confer antibiotic resistance can arise following evolution in LB broth in the absence of an antimicrobial, raising questions over whether evolutionary trajectories *in vitro* will be the same *in vivo* due to adaption to a specific environment [8]. Therefore, if *in vitro* studies are used to guide changes in treatment and prescription policy, it is important that the clinical community have the confidence in the real-world translation of the experimental observations.

Another recent study investigated whether compensatory mutations that arise following acquisition of AMR genes by *Escherichia coli* had the same effect in different media; the results showed that such compensation does not always translate in another environment [5]. This is an import finding, suggesting that a compensatory mutation can be beneficial in one clinical setting, such as the bladder, but may be deleterious in another, for example the blood [5]. It further highlights the importance of the choice of growth media when assessing fitness effects following acquisition of resistance.

In a recent study, the incidence of bacteraemia in England following an initial *Escherichia coli* infection of the urogenital tract was high (51.2%) [13], with a large number of these isolates found to be resistant to trimethoprim (48.1%), ciprofloxacin (18.5%) and amoxicillin-clavulanic acid (AMC) (45.4%,) [13]. Therefore, studying antimicrobial resistance in *Enterobacteriaceae* with better models is of particular interest as resistance continues to increase within this family to such a degree that carbapenem resistant, ESBL producing *Enterobacteriaceae* are now considered priority pathogens on the critical list, to which new drugs are urgently required [14].

There have been many *in vitro* models developed to study bacteria that more closely mimic their natural habitats within the human body; these include the chemostat gut, which has been used to study both planktonic and biofilm growth of *Clostridium difficile* [15, 16], and fermenter models of the human oral cavity, which have been used to study AMR and horizontal gene transfer [17, 18]. More relevant still are laboratory-grown organoids which allow pathogen-host interactions to be investigated and observed in isolation [19–21]. A recently described urothelial organoid was shown to be tolerant of urine over extended periods of time, to produce a number of expected biomarkers in the presence of urine, and to respond to bacterial infection in a fashion similar to that of the human bladder [21].

In this paper, we initially evaluated whether the type of media used during resistance development affects the evolutionary trajectory of the bacterial strain in the presence of sub-inhibitory concentrations of an antibiotic. We chose as a model *Escherichia coli* 10129, a fully susceptible clinical isolate from a patient with clinical suspicion of meningitis in Malawi [22], a country with high rates of bacteraemia caused by *E. coli* (69% of bacteraemia caused by *E. coli* in 2016), many of which are multidrug resistant [23], including to amoxicillin-clavulanic acid, which is commonly used. We included representatives of different media used in previous evolutionary studies, M9 (defined media) [6], iso-sensitest broth (ISO, semi-defined media) [24] and LB broth (LB, undefined) [8]. The whole genomes of six AMC-resistant derivatives of *E. coli* 10129 were sequenced to identify and predict mutations within the genome likely to account for observed AMC resistance. Using these six isolates, we next investigated the effect of media on the assessment of both minimum inhibitory concentrations (MIC) and the competitive fitness of the AMC-resistant derivatives. Finally, we examined whether MICs and fitness determined in growth media could be used to predict the MIC determined in the more physiological environment of urine, and comparative growth in urothelial organoids.

## Materials and methods

### Bacterial strains, media and antibiotics

The ancestor bacterial strain used in this study was the clinical isolate *E. coli* 10129 which was isolated from a cerebral spinal fluid (CSF) sample in Malawi [22] (Table 1). It was reported as fully susceptible and the genome had previously been resolved to a relatively small number of contigs within this strain collection [19].

**Table 1:** *Escherichia coli* isolates used during this study

| ***Escherichia coli* Isolate** | **Comments** | **Origin** |
| --- | --- | --- |
| 10129 | Sensitive clinical Malawian isolate. The ancestral strain used for this study. | Musicha *et al.* 2017 [22] |
| ISO_2 | AMC-resistant derivative of *E. coli* 10129 selected for in Iso-sensitest broth | This study |
| ISO_9 | AMC-resistant derivative of *E. coli* 10129 selected for in Iso-sensitest broth | This study |
| ISO_10 | AMC-resistant derivative of *E. coli* 10129 selected for in Iso-sensitest broth | This study |
| 10129 LB_1 | AMC-resistant derivative of *E. coli* 10129 selected for in LB broth | This study |
| 10129 LB_2 | AMC-resistant derivative of *E. coli* 10129 selected for in in LB broth | This study |
| 10129 LB_5 | AMC-resistant derivative of *E. coli* 10129 selected for in in LB broth | This study |

Cultures were grown in Mueller Hinton broth (MHB), LB (Lennox) (both Sigma, UK), Isosensitest (ISO) (Oxoid, UK) or M9 (50% (v/v) M9 minimal salts (2x) (Gibco, ThermoFisher Scientific, USA), 0.4% D-glucose, 4mM magnesium sulphate (both Sigma, UK) and 0.05 mM calcium chloride (Millipore, USA)). *E. coli* 10129 and AMC-resistant isolates were grown on LB agar (Lennox) (Sigma, UK) for colony counting during MIC determination.

Amoxicillin trihydrate: potassium clavulanate (4:1) (AMC) was diluted in molecular grade water (both Sigma, UK) to a stock concentration of 1 mg/ml and filter sterilised through a 0.22 μM polyethersulfone filter unit (Millipore, USA).

### Determination of minimum inhibitory concentration in laboratory growth media

MICs were performed in MHB, LB, ISO or M9, using cultures grown in the respective media in the absence of antimicrobial selection, following the Clinical and Laboratory Standards Institute (CLSI) guidelines for broth microdilution MIC and determined visually [25].

### Determination of minimum inhibitory concentrations in urine

Overnight bacterial cultures of *E. coli* 10129 and AMC-resistant isolates were diluted in urine supernatants derived from the organoids, with the composition of approximately 25% urine and 75% CnT-Prime 3D Barrier medium (CnT-PR-3D) (CELLnTEC, Switzerland), to OD_600_ of 0.1. Cultures were further diluted 1/1000. Antimicrobial susceptibility was determined by following the CLSI guidelines. Optical density (OD_595_) was measured at 24 h post-incubation with a microplate reader (Biochrom EZ Read 400).

### Selection of amoxicillin-clavulanic acid resistant E. coli 10129

*E. coli* 10129 was grown in sub-inhibitory concentrations (4 μg/ml), as determined by CLSI guidelines in MHB, of AMC in LB, ISO and M9 to select for resistant isolates. Separate cultures of *E. coli* 10129 were initially grown in 10 ml LB, ISO or M9 for 18 hours at 37°C and shaking at 200 rpm. Following incubation, each culture was serially diluted 1 in 10 in phosphate buffered solution (PBS, pH 7.2, Gibco, ThermoFisher Scientific, USA) and 50 μl of the neat to 10^−7^ dilutions were plated out on to LB agar and incubated at 37°C for 18 hours to determine CFU/ml. Using the initial cultures, 10 μl was diluted in 10 ml of the respective test media containing 4 μg/ml AMC and incubated at 37°C with shaking at 200 rpm for 24 hours. The cultures were serially diluted in PBS as described above and 50 μl of the neat to 10^−7^ dilutions were plated onto LB agar and 50 μl of neat to 10^−2^ dilutions were plated onto LB agar + 8 μg/ml AMC (MIC) and LB agar + 16 μg/ml AMC (2 x MIC). These plates were incubated at 37°C for 18 hours. Up to 10 single colonies were selected from agar plates from three replicate resistance selection experiments which had growth at the highest concentration of AMC and immediately stored in 1 ml LB + 40% glycerol (Sigma, UK) and stored at −80°C to prevent any further growth and compensatory mutations. The number of colonies was scored at the dilution that grew between 15-100 colonies, except for those that only grew a small number of colonies in the undiluted sample.

### Competitive fitness assays

The competitive fitness of each of the selected AMC-resistant isolates was determined in comparison to the ancestral isolate, *E. coli* 10129. Cultures of ancestor and AMC-resistant isolates were initially inoculated in LB, ISO or M9 directly from the −80°C stocks to minimise any further evolutionary events. The ancestor isolate was grown without the presence of AMC, whereas the resistant isolates were grown in the presence of the same concentration of AMC in which they were initially derived to confirm resistance and ensure that only the resistant population grew, i.e. 8 or 16 μg/ml AMC. All cultures were diluted in the appropriate media to an optical density at 600nm (OD_600_) of 0.1. The AMC-resistant and ancestor isolates were further diluted 1/1000 and combined 1:1 in the same media, 150 μl of which was added to a 96 well plate and incubated at 37°C for 24 hours at 200 rpm. The initial combined culture was serially diluted in PBS and 50 μl of the neat to 10 ^-7^ dilutions were plated out onto both LB agar and LB agar plus either 8 or 16 μg/ml AMC, depending on the concentration of AMC that the isolate was selected on, and incubated at 37°C for 18 hours. Following 24 hours of growth, the combined culture was then plated out on to agar as described above. The number of colonies was scored at the dilution that grew between 15-120 colonies.

Relative fitness was calculated using the Malthusian equation [26]:

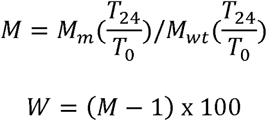

where *W* is the relative fitness, *M* is the Malthusian parameter, *M*_*wt*_ is the Malthusian parameter of the ancestor isolate, *M*_*m*_ is the Malthusian parameter of the resistant isolate, *T*_*0*_ is the log CFU/ml count of the initial culture and *T*_*24*_ is the log CFU/ml count after 24 hours.

### DNA extraction

AMC-resistant isolates and the ancestral isolate were grown in 10 ml LB and incubated at 37°C with shaking at 200 rpm for 18 hours. DNA extraction was performed using the Gentra Puregene Yeast/Bact. Kit (Qiagen, Germany) according to manufacturer’s instructions. The pellet was air-dried at room temperature for 5 minutes. The DNA was dissolved in 50 μl molecular grade water for 1 hour at 65°C and the DNA concentration determined using a NanoDrop™ One/One^c^ Microvolume UV-vis Spectrophotometer (ThermoFisher Scientific, USA).

### Comparative growth in bladder organoids

Human bladder urothelial organoids were prepared as described previously [21]. Briefly, primary human bladder epithelial cell (HBLAK) (CELLnTEC, Switzerland) were seeded onto polycarbonate transwell inserts (VWR, UK) at 5 × 10^5^ cells per ml in CnT-Prime medium (CnT-PR, CELLnTEC, Switzerland). An appropriate amount of CnT-PR was added to the basolateral chamber of the inserts so that the medium levels were equal, and the cells were submerged. Once cells reached confluency, CnT-PR was replaced with CnT-PR-3D (CELLnTEC, Switzerland) and incubated for 15-16 h. 3D culture was initiated by replacing medium from the apical chamber with filter-sterilised human urine (BIOIVT, UK), as well as fresh basolateral CnT-PR-3D medium. Urine and CnT-PR-3D medium were changed at regular intervals and organoids were fully established and stratified on day 14-18.

All bacterial cultures of *E. coli* 10129 and AMC-resistant isolates were diluted in urine to OD_600_ of 0.1, then further diluted 1/1000. Urine from apical chambers were aspirated and replaced with diluted bacterial cultures and fresh CnT-PR-3D was added to the basolateral chambers. Inserts were incubated at 37 °C, 5% CO_2_ for 24 h. Cultures from apical chambers were collected and transferred into a 96-well plate. Optical density (OD_595_) was measured at time 0 and 24 h post-infection using a microplate reader (Biochrom EZ Read 400).

Background optical density at 595 nm was accounted for by normalising with a measurement of the bladder organoids containing the *E. coli* 10129 ancestor or AMC-resistant isolates at 0 hours, as well as for organoid containing media only. Relative growth was calculated using the following equation:

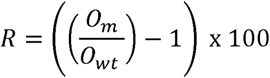

where *R* is the relative growth, *O*_*wt*_ is the OD595 after 24 hours growth of the AMC-resistant isolate in the bladder organoid, *O*_*m*_ is the OD595 after 24 hours growth of the ancestor isolate in the bladder organoid.

### Whole genome sequencing and bioinformatic analysis

Whole genome sequencing of *E. coli* 10129 and AMC-resistant isolates were performed by MicrobesNG (MicrobesNG, UK), which also included the trimming of the sequencing reads. *De novo* assembly of each of the genomes were performed using SPAdes (version 3.12.0) [27] and assembly statistics were generated using QUAST (version 4.6.3) [28]. Annotation of each of the assembled genomes was performed using Prokka (version 1.12) [29]. Single nucleotide polymorphisms (SNPs) in each of the *E. coli* 10129 AMC-resistant isolates were predicated computationally using Breseq (version 0.32.0) in default mode, with the trimmed sequencing reads produced by MicrobesNG aligned against our assembled and annotated genome of the *E. coli* 10129 ancestor isolate. Any SNPs which were predicted with less than 80% frequency were discounted. Also discounted were stretches of missing coverage which mapped to small, full contigs from the ancestor and a single 192 bp duplication at the end of a contig from LB_5 which was deemed an assembly artefact (Table S1).

### PCR confirmation of predicted single nucleotide polymorphisms

Primers were designed to amplify a 465bp region of the *cpxA* gene containing the predicted SNP and a 442bp region that contained the predicted SNP within the *ampC* promoter region (Table 2). The PCR amplification was performed using the Q5^®^ High Fidelity polymerase (New England Biolabs, USA) and the final reaction contained lx Q5^®^ reaction buffer, 200 μM dNTPs, 0.5 μM of the appropriate forward and reverse primers listed in Table 2 and 0.02 U/μl Q5^®^ polymerase in a total volume of 25 μl. The PCR samples were run using the following protocol: denaturation at 98°C for 30 seconds, followed by 35 cycles of denaturation at 98°C for 10 seconds, annealing at 69°C (*cpxA* SNP) or 67°C (*ampC* SNP) for 30 seconds, elongation at 72°C for 20 seconds, followed by a final extension of 2 minutes at 72°C. The PCR products were cleaned up using the Monarch^®^ PCR and DNA clean-up kit (New England Biolabs, USA). The PCR samples were mixed with 125 μl of the DNA Clean-Up Binding Buffer, transferred to the Clean-Up Columns and centrifuged at 11400 *x g* for 1 minute. The bound DNA was washed with 200 μl DNA Wash Buffer and centrifuged at 11400 *x g* for 1 minute, twice, followed by elution in 20 μl molecular grade water (Sigma, UK) by centrifuging at 11400 *x g* for 1 minute after incubating at room temperature for 2 minutes. All PCR products were Sanger sequenced at GeneWiz (Takely, UK).

**Table 2:**
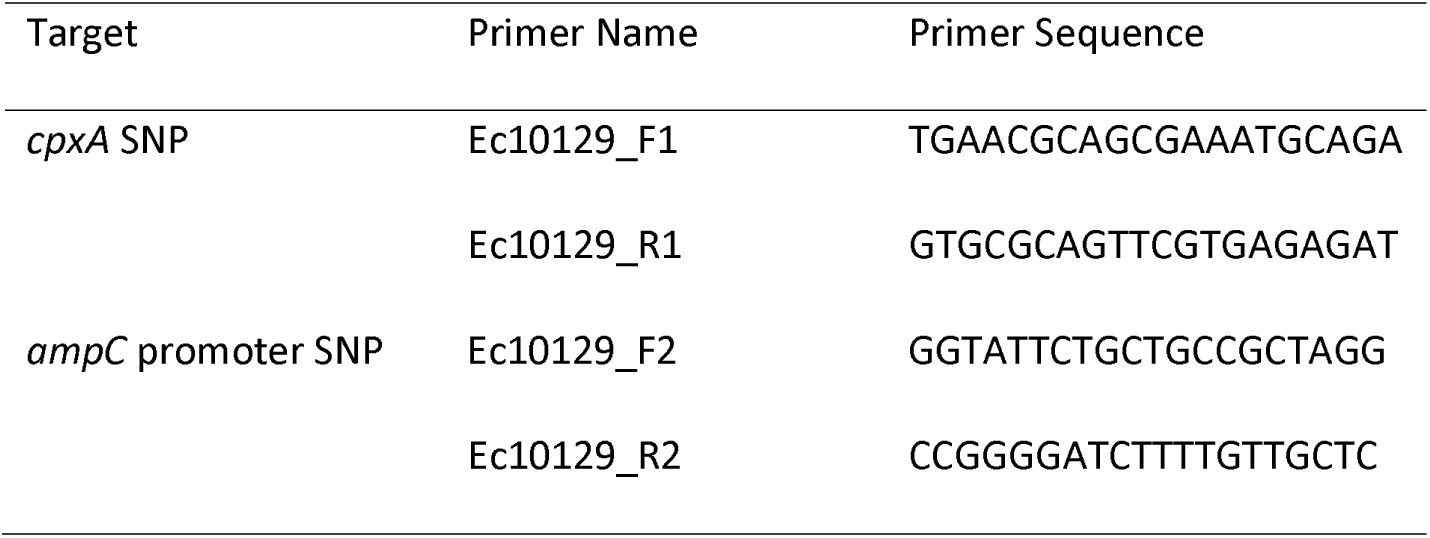
Primer names and primer sequences for confirmation of the SNPs in the *cpxA* gene and *ampC* promoter region of *E. coli* 10129 ancestor and AMC-resistant isolates as predicted by breseq

### Biofilm assay

*E. coli* 10129 ancestor and AMC-resistant isolates were grown in LB for 18 hours at 37°C and 200 rpm and then diluted 1/1000 in M9. Three 100 μl technical replicates of the each of the diluted cultures were added to a 96 well microtitre plate alongside three technical replicates of M9 only and three empty wells and incubated statically at 37°C for 24 hours. Following incubation, all culture or media was removed, and the wells washed 4-5 times with 150 μl PBS and then left to dry upside-down for 10 minutes. The wells were then stained with 125 μl 0.1% Gram’s crystal violet solution (Sigma, UK) for 15 minutes at room temperature, then washed 4-5 times with 150 μl PBS and left to dry upside-down for between 60-90 minutes. The remaining stain was dissolved with 125 μl 30% acetic acid (Sigma, UK) and incubated at room temperature for 15 minutes. Finally, the acetic acid solution was transferred to a fresh 96 well microtitre plate and measured at an optical density of 550nm, with 30% acetic acid used as a blank and the stained empty wells as background.

### Statistical analysis

Statistical analysis of the initial growth of *E. coli* 10129 in LB, ISO and M9 was performed using Ordinary One-Way ANOVA in GraphPad Prism (version 8.0.0). Statistical analysis of competitive fitness in LB, relative fitness in bladder organoids and biofilm production of the *E. coli* 10129 ancestor and AMC-resistant isolates was performed using Ordinary One-Way ANOVA plus uncorrected Fisher’s LSD test in GraphPad Prism. Statistical analysis of competitive fitness of LB_2 and LB_5 in LB, ISO and M9 was performed using both Ordinary One-Way ANOVA plus uncorrected Fisher’s LSD test and 2-way ANOVA plus uncorrected Fisher’s LSD test. Finally, statistical analysis of cell density of *E. coli* 10129 following challenge with AMC in LB, ISO and M9 was performed using 2-way ANOVA plus uncorrected Fisher’s LSD test, also in GraphPad Prism.

## Results

### Selection of E. coli 10129 resistant isolates

The MIC of AMC for *E*. *coli* 10129 was determined to be 4-8 μg/ml (Table 3). To select for isolates showing decreased susceptibility to AMC, *E*. *coli* 10129 was grown in the presence of sub-inhibitory concentrations (4 μg/ml) of AMC in defined (M9), semi-defined (ISO) and undefined (LB) media for 24 hours.

**Table 3:** Minimum inhibitory concentrations of the *E. coli* 10129 ancestor and AMC-resistant isolates assessed in MHB, LB, ISO, M9 and urine following the CLSI guidelines.

| Medium | *E. coli* 10129 | ISO_2 | ISO_9 | ISO_10 | LB_1 | LB_2 | LB_5 |
| --- | --- | --- | --- | --- | --- | --- | --- |
| MHB | 4-8 µg/ml | 16 µg/ml | 8-16 µg/ml | 8-16 µg/ml | 32 µg/ml | 64 µg/ml | 64 µg/ml |
| LB | 4-8 µg/ml | 16 µg/ml | 8-16 µg/ml | 8-16 µg/ml | 32 µg/ml | 64-128 µg/ml | 64 µg/ml |
| ISO | 8 µg/ml | 16-32 µg/ml | 16 µg/ml | 16 µg/ml | 32 µg/ml | 128 µg/ml | 64-128 µg/ml |
| M9 | 1-2 µg/ml | 2-4 µg/ml | 2 µg/ml | 2 µg/ml | 4-8 µg/ml | 32 µg/ml | 16 µg/ml |
| Urine | 2 µg/ml | 4 µg/ml | 4 µg/ml | 4 µg/ml | 16 µg/ml | 64 µg/ml | 64 µg/ml |

There was no significant difference in the cell densities following growth of *E*. *coli* 10129 in the absence of AMC among three different media types (P value = 0.1071, Fig. 1a) and therefore the cell density of the inoculum was also comparable. A significant reduction in cell density of 5.049 log CFU/ml was observed following incubation in M9 containing AMC (P value = <0.0001, Fig. 1b) and we did not recover any isolates from agar containing 8 or 16 μg/ml AMC. Unlike M9, following exposure of *E*. *coli* 10129 to AMC in ISO there was a small but insignificant increase in cell density of 0.592 log CFU/ml (P value = 0.1339, Fig. 1b) and significant growth of *E. coli* 10129 of 1.03 log CFU/ml in LB in the presence of AMC compared with the initial inoculum (P value = 0.0235). Therefore, using this one-step selection technique, 10 colonies were selected from agar plates containing 8 μg/ml AMC and 10 from plates containing 16 μg/ml AMC following growth in ISO and LB broth, respectively.

**Figure 1:**
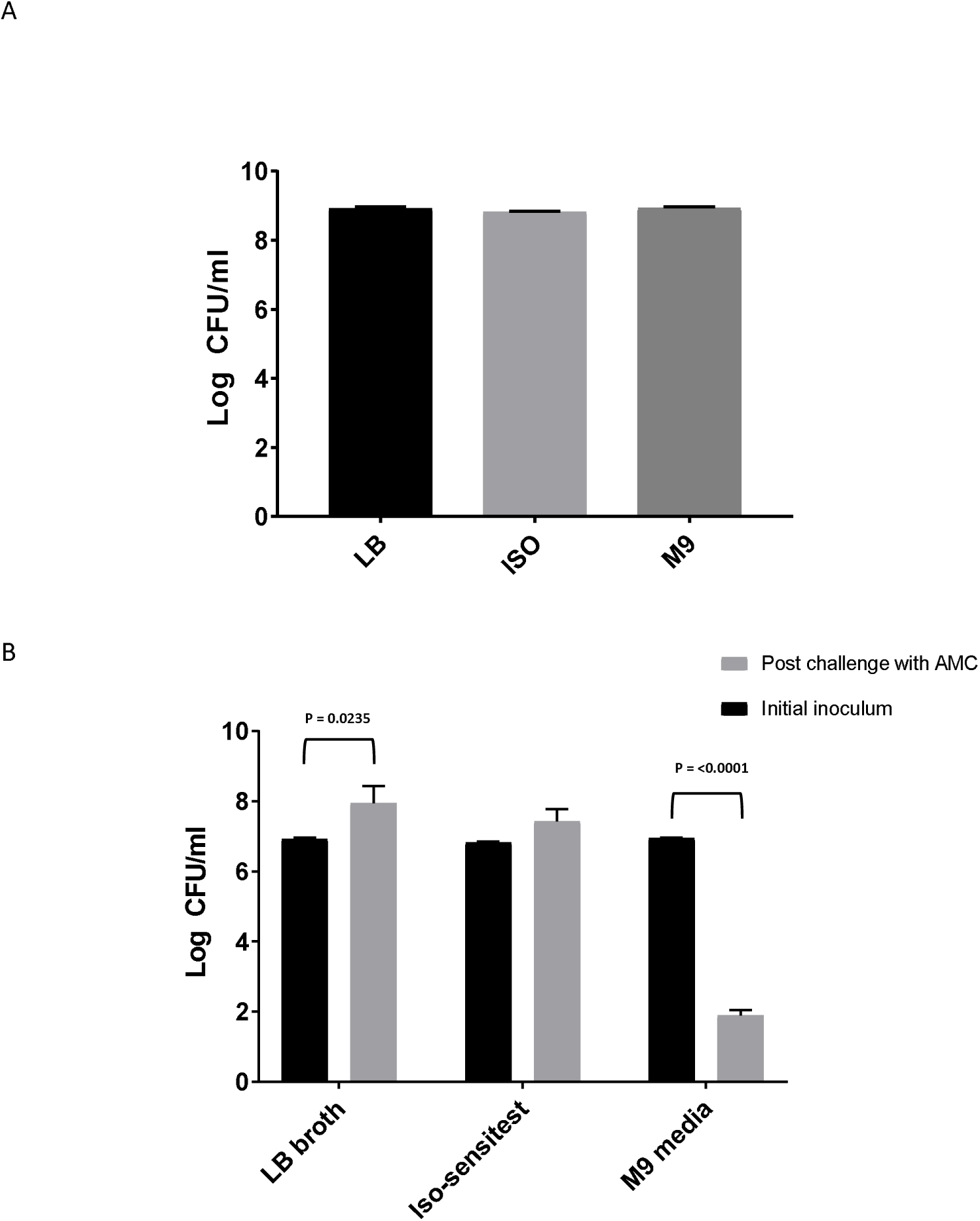
Difference in log CFU/ml of *E. coli* 10129 **(A)** after growth in LB, ISO and M9 and **(B)** following 24-hour exposure with sub-inhibitory concentrations of AMC in LB, ISO and M9 compared to the initial inoculum. Error bars represent standard error of the mean.

Of the 10 resistant isolates of *E. coli* 10129 grown in either LB or ISO, three from each media-type were selected at random. The three isolates selected following growth in ISO broth were designated ISO_2, ISO_9 and ISO_10 and those grown in LB were designated LB_1, LB_2 and LB_5.

### Minimum inhibitory concentrations of E. coli 10129 ancestor and AMC-resistant isolates in MHB

Resistance to AMC in the six selected AMC-resistant isolates were confirmed through MIC determination in comparison to the ancestral isolate in MHB (Table 3), and all were above the clinical breakpoints according to the EUCAST guidelines for systemic infections caused by *E*. *coli* (>8 μg/ml, Fig. 3). While there was within media-type increased variability in the AMC MIC of the resistant isolates, there was a clear distinction between those resistant isolates selected for in LB and those from ISO (Table 3). There was only a 1 to 3-fold increase in the AMC MIC in resistant isolates selected in ISO, which would be expected as ISO selected isolates where only recovered from agar containing MIC concentrations of AMC. The AMC-resistant isolates selected for in LB broth plated out on LB agar containing 2xMIC concentrations of AMC were all highly resistant compared with the ancestor isolate, resulting in a 3 to 15-fold increase in MIC (Table 3). The most resistant isolates were determined to be LB_2 and LB_5, both with a 7 to 15-fold increase in MIC in comparison to the ancestral isolate (Table 3). This data is summarised in Figure 3.

### Single nucleotide polymorphism prediction

SNPs that arose in the genomes of the *E. coli* 10129 AMC-resistant isolates following selection in sub-inhibitory concentrations of AMC were identified using Breseq, which predicts SNPs in genomes of derivative generations by mapping sequencing reads to the original ancestral isolate [30].

Chromosomal mutations of *E. coli* that lead to resistance to β-lactams are often found within the promoter region of the *ampC* gene, resulting in over-production of the chromosomally located β-lactamase AmpC [31–33]. Isolates LB_2 and LB_5 contained a predicted SNP at nucleotide position - 32 in the −35-box promoter region with 100% frequency within the sequencing reads, which was subsequently confirmed to be present using PCR and sequencing. This mutation converted the natural weak −35 promoter (TTGTCA) to a stronger promoter (TTGACA) [34] which has been previously linked to 21-fold increase in AmpC production [35].

A second SNP with a 100% frequency was predicted in isolate ISO_2, although there was only a modest increase in AMC MIC when assessed in MHB. This SNP occurred in *cpxA*, which encodes a change in amino acid from a proline to leucine at position 177. CpxA is the sensor histidine kinase portion of the envelope stress response two-component system, CpxAR. The CpxAR two-component system is involved in the regulation of several proteins including the efflux pumps *acrB, acrD*, and *eefB* [36] and the outer membrane porins *ompF* and *ompC* [37] in a wide range of Enterobacteriaceae including *E. coli.* Therefore, the SNP predicted by Breseq, and subsequently confirmed via PCR and sequencing, present in the *cpxA* gene may explain the modest increase in AMC resistance in ISO_2.

In LB_5, two synonymous SNPs were predicted with 80-90% frequency in the *vgrG1* gene, which is part of the type VI secretion system of Gram negative bacteria and a significant component for the delivery of effector molecules to other cells [38]. One SNP was found in *vgrG1* which mapped to amino acid position 368 (proline) and a second at amino acid 387 (glycine) and the same SNP, and indeed the only predicted SNP, at amino acid position 368 was also present in AMC-resistant isolate in LB_1 (Table S1).

There were no predicted SNPs within genes in AMC-resistant isolates ISO_9 or ISO_10, which suggests the modest increase in resistance may be due to tolerance mediated by increased gene expression or heteroresistance [39]. This has previously been seen in the overexpression of the resistance gene activator MarA in *E. coli* resulting in an increase in resistance to, and survival in, carbenicillin [40]. However, it is also possible that a mutation resulting in resistance was been missed due to gaps in short read sequencing.

### Competitive fitness of AMC-resistant isolates in LB

The relative fitness of the *E. coli* 10129 AMC-resistant isolates was assessed competitively in LB. There was a significant increase in fitness (16.6%) of LB_5 relative to the ancestral isolate (P value = 0.0283) and a significant increase in fitness (19.9%) relative to the ancestral isolate for ISO_2 (P value = 0.0122) (Fig 2A). The range of fitness effects observed between the three isolates selected for in LB (10.3% to 16.6%) was considerably smaller than range of fitness effects for the three isolates selected for in ISO (−3.4% to 19.9%). Therefore, we were able to produce AMC-resistant isolates with a range of fitness using the one-step selection technique.

**Figure 2:**
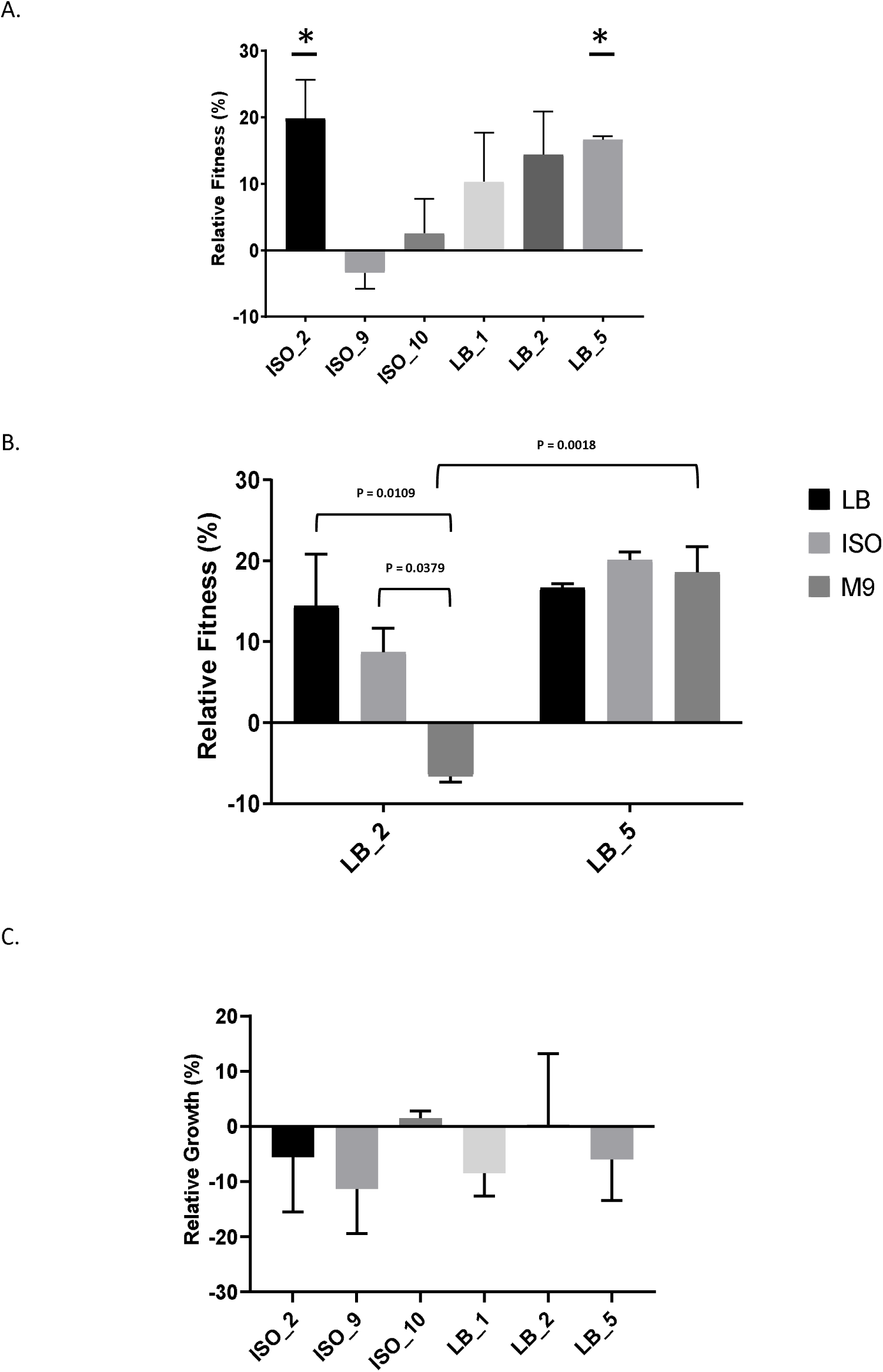
Relative fitness and growth of *E. coli* isolates; error bars represent standard error of the mean **A;** *E*. *coli* 10129 AMC-resistant isolates selected following growth in sub-inhibitory concentrations of AMC in either ISO or LB compared to the ancestral isolate in LB in the absence of AMC. **B;** Relative fitness of *E. coli* 10129 LB_2 and LB_5 AMC-resistant isolates in LB, ISO and M9, compared to the ancestral isolate in the absence of AMC. **C;** Relative growth of *E*. **coli** 10129 AMC-resistant isolates selected following growth in sub-inhibitory concentrations of AMC in either ISO or LB in comparison to the ancestral isolate in bladder organoids in the absence of AMC.

**Figure 3:**
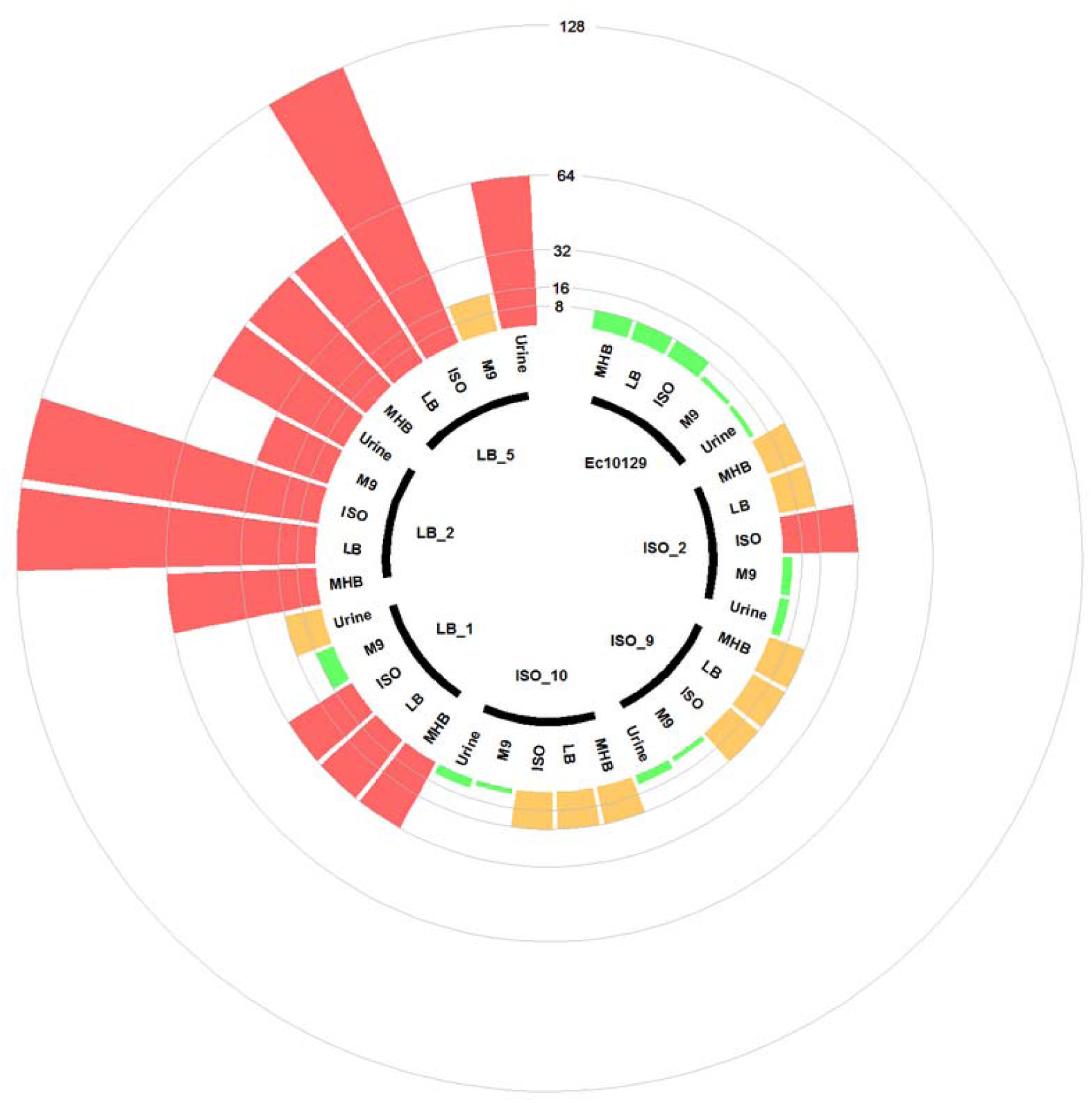
Circular barplot chart displaying the MIC values of the *E. coli* 10129 ancestor with 6 evolved derivative strains. The inner circle shows which bars belong to which strains and which media the MICs were tested. Clinical breakpoint for sensitivity and resistance are 8 μg/ml and 16 μg/ml respectively and are indicated by a circle. The scale represents the MIC value.

### Minimum inhibitory concentrations of E. coli 10129 ancestor and AMC-resistant isolates in other media and urine

Following observed differences between media types in the selection of AMC-resistant isolates *of E. coli* 10129, we assessed the MIC of the ancestor and AMC-resistant isolates in LB, ISO and M9 to determine whether media affects antimicrobial resistance (Fig. 3). MICs of both the ancestor and AMC-resistant isolates were comparable when assessed in MHB and LB; however, there was a slight increase in MIC in the ancestor isolate and ISO_2, LB_2 and LB_5 when assessed in ISO (Table 3, Fig. 3). There was a considerable decrease in MIC of all isolates when assessed in M9 compared with MHB, LB and ISO (Table 3, Fig. 3), which means the MIC of AMC in M9 (1-2 μg/ml) is actually lower than the sub-inhibitory concentrations of AMC used to select AMR resistant isolates in M9 (Table 3, Fig. 3). Therefore, the concentration of AMC used to select AMC-resistant isolates in M9 was not sub-inhibitory but rather bactericidal, explaining the observed decrease in cell density after exposure to AMC in M9 (Fig. 1B).

As the translation of laboratory results to the clinic requires more sophisticated models of growth environments than an agar plate or tube of media, we wanted to ascertain if the variability we see in the growth of the AMC resistant *E. coli* isolates described above is mirrored if they are grown in urine and with organoids. *E. coli* is the top urinary pathogen, responsible for upwards of 80% of all community-acquired urinary tract infections in otherwise healthy young women [41]. We therefore wanted to assess the performance of *E. coli* 10129 and its derivatives in the urinary environment.

We found that although the growth of *E. coli* 10129 was the same or better than that of the highly virulent uropathogenic *E. coli* strain UTI89 [42] in 100% urine (data not shown), pure urine is nutrient-poor and overall growth was not robust enough to determine MIC. We therefore used supernatant derived from the apical chamber of urothelial organoids, which consist of urine enriched with cellular exudate and which is more likely to be representative of the host/pathogen interface in the bladder [21]. MICs of the ancestor isolate, and the AMC-resistant isolates selected in ISO, assessed in this urine were noticeably lower than those assessed in MHB, LB and ISO and were more comparable to those assessed in M9 (Table 3, Fig. 3). The MIC of the AMC-resistant isolates selected for in LB assessed in urine was markedly higher than those assessed in M9 and are in fact more comparable to the MICs assessed in MHB. Taken together, these data suggest that culture environment can have an unpredictable effect on MIC.

### Competitive fitness of AMC-resistant isolates in LB, ISO and M9

As we have been able to show that media has an important effect on both the selection of *E. coli* 10129 AMC-resistant isolates and the MIC of the ancestor and resistant isolates, we determined the relative fitness of the highly resistant and fit LB_2 and LB_5 isolates in three media types in the absence of AMC: LB, ISO and M9. There was no significant difference in relative fitness when LB_5 was assessed in LB, ISO and M9 (P value = 0.4943) and no significant difference between the relative fitness of LB_2 and LB_5 when compared in LB (P value = 0.6548) or ISO (P value = 0.0533) (Fig. 2B). However, the relative fitness of LB_2 was significantly different when assessed in M9 media compared with its growth in ISO (P value = 0.0379) and LB (P value = 0.0109) (Fig. 2B). Additionally, there was a significant difference in relative fitness between LB_2 and LB_5 when assessed in M9 media (P value = 0.0018) (Fig.2B). These data suggest that media choice can also affect fitness.

### Comparative growth rates in bladder organoids

As many studies into the evolution or acquisition of AMR and subsequent effects on fitness are ultimately intended to be translated into clinical interventions, we determined whether relative fitness in LB can be used to predict relative growth of the *E. coli* 10129 AMC-resistant isolates compared with the ancestral isolate in urothelial organoids. Despite relative growth varying among the AMC-resistant isolates, no significant difference in relative growth among all six AMC-resistant isolates was observed (P value = 0.8592) (Fig. 2C) and, in direct contrast to relative fitness assessed in LB, most AMC-resistant isolates grew more slowly relative to the ancestral isolate. The relative growth of ISO_2 was 5.6% slower compared to the ancestral isolate despite previously being found to have acquired the largest increase in fitness (19.9%) of the six AMC-resistant isolates in LB (Fig. 2C). Although LB_1 and LB_5 increased in fitness in LB (10.3% and 16.6%, respectively), and in the case of LB_5 an increase in fitness in ISO and M9 as well, both isolates grew more slowly (8.5% and 6.1% slower, respectively) relative to the ancestral isolate in the bladder organoid (Fig. 2C). Isolate ISO_10’s fitness increase in LB (2.5%) was comparable to the relative growth in the bladder organoid (1.5%). This indicates that relative fitness assessed in growth media, and in particular LB, has no predictive value to the relative growth in more biologically relevant environments.

### Biofilm production

Urinary tract infection (UTI) isolates often form extensive biofilms on catheter surfaces and we wanted to determine whether the fitness of *E. coli* 10129 AMC-resistant isolates was accompanied by an increase in biofilm production. While biofilm production varied in the AMC-resistant isolates, there was an increase in biofilm production for all AMC-resistant isolates in comparison to the ancestor isolate except for LB_5, which was previously found to acquire a fitness benefit. Also, despite ISO_9 incurring a fitness cost in LB and growing more slowly compared to ancestral isolate in bladder organoids, there was no significant difference in biofilm production compared with the ancestral isolate (P value = 0.3532) (Fig. 4). There was a significant increase in biofilm production compared with the ancestor isolate in only two isolates; LB_1 (3.97-fold increase, P value = <0.0001) and LB_2 (1.90-fold increase, P value = 0.0147) (Fig. 4), even though LB_1 grew more slowly (8.45%) and LB_2 marginally grew faster (0.331%) than the ancestral isolate in bladder organoids. Therefore, we found no correlation between the fitness of AMC-resistant isolates in different growth conditions and biofilm production.

**Figure 4:**
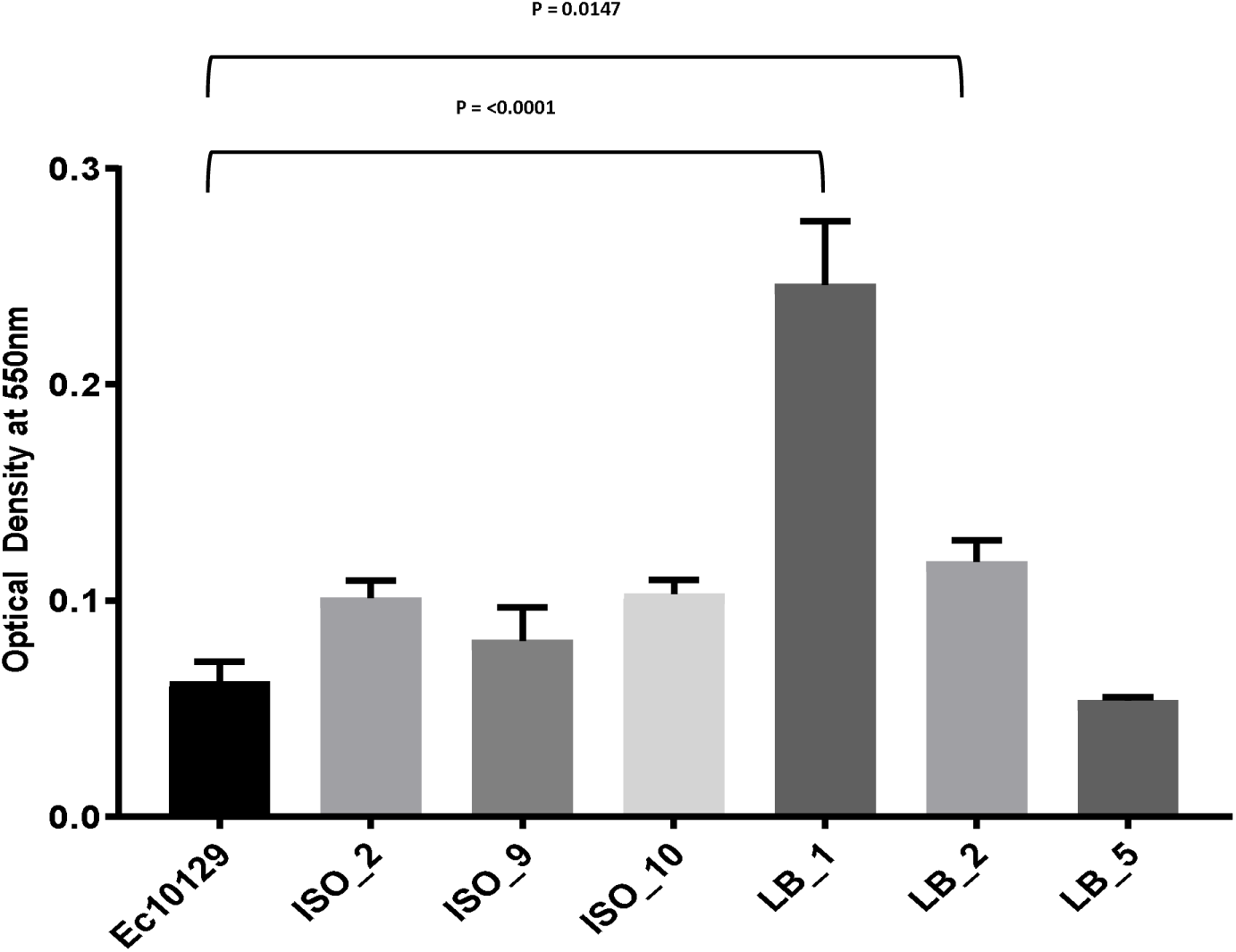
Biofilm production of *E. coli* 10129 ancestor and AMC-resistant isolates in M9. Error bars represent standard error of the mean

## Discussion

The prospect of using reproducible evolutionary trajectories to resistance as a tool to control the emergence and persistence of AMR is a growing area of research [3, 4, 43]. One of the prerequisites for translation of these ideas to the clinic in the form of informed prescription policy advice is robust and reproducible data which can convince clinicians and policy makers that what is seen in the laboratory will actually occur in patients and the wider healthcare environment. Therefore, there is a need to analyse the consequences of different experimental conditions thoroughly and ensure predictive value as studies move into the clinic.

This study is part of a continuing One Health investigation into the drivers of AMR in Malawi which has a high rate, and increasing incidence, of antimicrobial resistance [23]. Between 1998 and 2016, 66% of bacteraemia caused by *E. coli* in Malawi were found to be multidrug resistant, and the incidence of resistance to second-line antibiotics is on the increase [23]. To generate resistant isolates in the laboratory, we used a fully susceptible Malawian clinical isolate *E. coli* 10129 [22], and exposed it to AMC *in vitro* as this is a geographically and clinically relevant first line treatment for Enterobacteriaceae infections [23]. We used a single-step procedure in three different media types that are commonly used in evolutionary studies [6, 8, 24] to select for AMC-resistant *E. coli* 10129 isolates with AMC-resistance which were then tested for relative fitness compared with the ancestral strain [5, 8, 24, 26]. We were able to determine multiple mutations leading to AMC resistance such as well-characterised SNPs in the *ampC* promoter region [31–35], and others with less well defined evidence of AMC resistance including SNPs in *cpxA* or *vrgG* [44–46].

Several separate SNPs in the *cpxA* gene have previously been linked to aminoglycoside [44] and β-lactam [45] resistance following *in vitro* selection. These previously reported SNPs resulted in non-synonymous amino acid changes at the follow positions of Leu-57-Val [45], Ala-183-Gly, Ile-95-Phe, Met-22-Arg and Trp-184-Gly [44] whereas the non-synonymous SNP we identified within *cpxA* in *E. coli* 10129 was an amino acid change of Pro-Leu at position 177.

Recently a knockout of *vgrG* in *Acinetobacter baumannii* has been linked to ampicillin-sulbactam resistance, although AMC resistance due to *vgrG* knockout could not be determined due to the strain used already having high-level resistance to AMC [46].

In a clinical setting, MICs are standardised and are determined in MHB/cation-adjusted MHB according to either CLSI [25] or EUCAST [47] guidelines. Various evolutionary studies assess MICs in the same media that is subsequently used for selection of spontaneous AMR mutants, such as M9, ISO and LB [5, 6, 24]. However, other studies occasionally one medium, such as ISO [48] or MHB [49], to perform MICs and a different medium to perform subsequent evolutionary experiments, including nutrient broth [48] and LB [49]. We found that media can have a direct effect on the MIC, most notably an obvious drop in MIC when assessed in M9 compared to MHB, LB and ISO, and there were also more subtle differences between the latter three media. A recent study has found that supplements added to MHB, such as human serum or lung surfactant, directly affected the activity of antibiotics and therefore the determination of the MIC [50]. While we understand that the use of MHB in a clinical setting is entirely to ensure standardisation across different clinical laboratories to be able to monitor AMR, there is currently no comparable standardisation to use for evolutionary studies. To be able to compare studies from different laboratories and allow the cumulative picture of the evolutionary landscape of resistance development in pathogens to form, it is fundamental that these studies be comparable in terms of growth media and conditions used.

We have found that media not only has a direct effect on the selection of spontaneous mutants conferring resistance to AMC in our experiments, but it also affects the competitive fitness of AMC-resistant strains selected for in LB. Although both LB_2 and LB_5 contain the same mutation in the *ampC* promoter region which conferred resistance to AMC [34, 35], it is likely that either LB_5 has a compensatory mutation that allows it to grow better in M9 or there is a mutation elsewhere within the genome of the strains which results in a negative (for LB_2) or positive epistatic (for LB_5) interaction(s) only detectable during growth in M9 (Table S1). It is noteworthy that any compensatory mutations or mutations resulting in epistatic interactions identified in the genome would not be able to be linked to the growth phenotype unless comparative fitness assays were carried out in the different media. A similar phenomenon has previously been observed when competitive fitness was assessed in LB, M9 and tryptone soya broth following adaption of *E.coli* K12 MG1655 to LB to select for mutations that compensate for a loss of fitness following a mutation in either *gyrA* or *marR* [5]. Both this study and ours highlights that mutations, and the associated fitness effects, can have different consequences in different environments.

To determine if any of the data we derived from growth in laboratory media could have predictive value in a more clinically relevant environment, we assessed growth in urine and urothelial organoids. We found that MICs of the *E. coli* 10129 AMC-resistant isolates, when determined in urine, were not directly comparable to any media type we tested in this study and, more importantly, that the relative fitness of the AMC-resistant isolates was not comparable to relative growth in bladder organoids. The human bladder organoid model system used in this study was deliberately chosen as it closely represents the stratified and differentiated bladder urothelium, which is a common, and clinically important, *in vivo* environment for *E. coli* [21]. The availability of cellular constituents secreted into the urine from the epithelial cells of the bladder organoid, as well as nutrients and attachment factors present on the urothelial surface itself, along with its elaboration of a glucosaminoglycan layer, are all likely to have a direct effect on bacterial growth, therefore affecting the growth and MIC of bacteria, neither of which would be accurately predicted by growth in laboratory media.

As biofilm-producing bacteria are a major cause of catheter-associated UTIs [51], of which *E. coli* is the amongst the most common cause [51], we sought to determine whether there is any association between biofilm formation, MICs and fitness assessed in the different growth media, including urine and urothelial organoids. We found no correlation between fitness, MICs and biofilm production in our *E. coli* 10129 derivative strains suggesting that this lack of predictive value between MIC and fitness in different media is also evident between MIC, fitness and biofilm forming ability for this strain.

We acknowledge there are limitations to this study, not least the limited amount of different *E. coli* isolates, bacterial strains and antimicrobials tested, and the limited number of derivative isolates tested following selection. Whilst we had a very limited number of fully susceptible clinical isolates from Malawi to choose from [22], it is likely that in the clinic only a single isolate will be responsible for infection. In addition, the use of multiple evolutionary lineages derived from a single strain is sufficient to demonstrate variability and reflects the evolving epidemiological landscape of *E. coli* UTIs [52, 53].

These results highlight the importance of not only assessing aspects of evolutionary studies in several media types but also the importance of using *in vitro* model systems, such as bladder organoids, to ensure the phenomenon that is observed occurs in a range of different settings rather than in a single environment. For example, a diagnostic laboratory using MHB to determine sensitivity of the ISO-derived strains in this study would have deemed them resistant to AMC, whereas the organoid result would report that the strains were actually sensitive. If the MHB-derived information were deployed in a clinical setting, this could lead to incorrect treatment of the patient, resorting to a less-optimal alternative or delaying the administration of a more suitable drug. Indeed, in our clinical experience with chronic UTI patients, mismatches between predicted and real-world response outcomes of antibiotic treatment are not uncommon; our results could in part explain these discrepancies.

## Conclusions

Media used in evolutionary studies to select for mutations or horizontal gene transfer events that result in resistance to antimicrobials can affect the outcome. We found that MICs and competitive fitness assessed in growth media do not predict or correlate directly to MICs or relative growth when determined in urine or urothelial organoids. In studies with the aim of informing antibiotic treatment regimens, particular attention should be given to the study design with respect to media and growth models used.

## Acknowledgements

We would like to thank the MRC (grant numbers MR/R015074/1 and MR/S004793/1) for funding.

## Accession numbers

This assembled genome of the ancestral isolate, *E. coli* 10129, has been deposited at GenBank under the accession SIJF00000000. The version described in this paper is version SIJF01000000. The sequencing reads of the six AMC resistant isolates derived from the ancestral isolate and submitted under the accession number PRJNA522956.

## References

1. Singer, A.C., et al., Review of Antimicrobial Resistance in the Environment and Its Relevance to Environmental Regulators. Front Microbiol, 2016. 7: p. 1728.

2. Caudell, M.A., et al., Identification of risk factors associated with carriage of resistant Escherichia coli in three culturally diverse ethnic groups in Tanzania: a biological and socioeconomic analysis. The Lancet Planetary Health, 2018. 2(11): p. e489–e497.

3. Wong, A., Epistasis and the Evolution of Antimicrobial Resistance. Front Microbiol, 2017. 8: p. 246.

4. Hughes, D. and D.I. Andersson, Evolutionary Trajectories to Antibiotic Resistance. Annu Rev Microbiol, 2017. 71: p. 579–596.

5. Basra, P., et al., Fitness tradeoffs of antibiotic resistance in extra-intestinal pathogenic Escherichia coli. Genome Biol Evol, 2018.

6. Suzuki, S., T. Horinouchi, and C. Furusawa, Acceleration and suppression of resistance development by antibiotic combinations. BMC Genomics, 2017. 18(1): p. 328.

7. Palmer, A.C., et al., Delayed commitment to evolutionary fate in antibiotic resistance fitness landscapes. Nat Commun, 2015. 6: p. 7385.

8. Knöppel, A., J. Näsvall, and D.I. Andersson, Evolution of Antibiotic Resistance without Antibiotic Exposure. Antimicrobial Agents and Chemotherapy, 2017. 61(11): p. e01495–17

9. Westhoff, S., et al., The evolution of no-cost resistance at sub-MIC concentrations of streptomycin in Streptomyces coelicolor. ISME J, 2017. 11(5): p. 1168–1178.

10. Gullberg, E., et al., Selection of resistant bacteria at very low antibiotic concentrations. PLoS Pathog, 2011. 7(7): p. e1002158.

11. Starikova, I., et al., Fitness costs of various mobile genetic elements in Enterococcus faecium and Enterococcus faecalis. J Antimicrob Chemother, 2013. 68(12): p. 2755–65.

12. Olivares Pacheco, J., et al., Metabolic Compensation of Fitness Costs Is a General Outcome for Antibiotic Resistant Pseudomonas aeruginosa Mutants Overexpressing Efflux Pumps. mBio, 2017. 8(4): p. e00500–17

13. Abernethy, J., et al., Epidemiology of Escherichia coli bacteraemia in England: results of an enhanced sentinel surveillance programme. J Hosp Infect, 2017. 95(4): p. 365–375.

14. World Health Organisation. WHO publishes list of bacteria for which new antibiotics are urgently needed. 2017. World Health Organisation. https://www.who.int/news-room/detail/27-02-2017-who-publishes-list-of-bacteria-for-which-new-antibiotics-are-urgently-needed.

15. Crowther, G.S., et al., Development and validation of a chemostat gut model to study both planktonic and biofilm modes of growth of Clostridium difficile and human microbiota. PLoS One, 2014. 9(2): p. e88396.

16. Baines, S.D., J. Freeman, and M.H. Wilcox, Effects of piperacillin/tazobactam on Clostridium difficile growth and toxin production in a human gut model. J Antimicrob Chemother, 2005. 55(6): p. 974–82.

17. Ready, D., et al., Composition and antibiotic resistance profile of microcosm dental plaques before and after exposure to tetracycline. Journal of Antimicrobial Chemotherapy, 2002. 49(5): p. 769–775.

18. Roberts, A.P., et al., Transfer of TN916-like elements in microcosm dental plaques. Antimicrob Agents Chemother, 2001. 45(10): p. 2943–6.

19. Forbester, J.L., et al., Interaction of Salmonella enterica Serovar Typhimurium with Intestinal Organoids Derived from Human Induced Pluripotent Stem Cells. Infect Immun, 2015. 83(7): p. 2926–34.

20. Leslie, J.L., et al., Persistence and toxin production by Clostridium difficile within human intestinal organoids result in disruption of epithelial paracellular barrier function. Infect Immun, 2015. 83(1): p. 138–45.

21. Horsley, H., et al., A urine-dependent human urothelial organoid offers a potential alternative to rodent models of infection. Sci Rep, 2018. 8(1): p. 1238.

22. Musicha, P., et al., Genomic landscape of extended-spectrum beta-lactamase resistance in Escherichia coli from an urban African setting. J Antimicrob Chemother, 2017. 72(6): p. 1602–1609.

23. Musicha, P., et al., Trends in antimicrobial resistance in bloodstream infection isolates at a large urban hospital in Malawi (1998–2016): a surveillance study. The Lancet Infectious Diseases, 2017. 17(10): p. 1042–1052.

24. Farrell, D.J., et al., Investigation of the potential for mutational resistance to XF-73, retapamulin, mupirocin, fusidic acid, daptomycin, and vancomycin in methicillin-resistant Staphylococcus aureus isolates during a 55-passage study. Antimicrob Agents Chemother, 2011. 55(3): p. 1177–81.

25. Clinical and Laboratory Standards Institute. Methods for Dilution Antimicrobial Susceptibility Tests for Bacteria That Grow Aerobically. 11th Ed. CLSI Standard M07. 2018 Clinical and Laboratory Standards Institute. Wayne, Pennsylvania, USA.

26. Lenski, R.E., et al., Long-Term Experimental Evolution in Escherichia coli: Adaptation and Divergence During 2,000 Generations. The American Naturalist, 1991. 138(6): p. 1315–1341.

27. Bankevich, A., et al., SPAdes: a new genome assembly algorithm and its applications to single-cell sequencing. J Comput Biol, 2012. 19(5): p. 455–77.

28. Mikheenko, A., et al., Versatile genome assembly evaluation with QUAST-LG. Bioinformatics, 2018. 34(13): p. i142–i150.

29. Seemann, T., Prokka: rapid prokaryotic genome annotation. Bioinformatics, 2014. 30(14): p. 2068–9.

30. Deatherage, D.E. and J.E. Barrick, Identification of mutations in laboratory-evolved microbes from next-generation sequencing data using breseq. Methods Mol Biol, 2014. 1151: p. 165–88.

31. Siu, L.K., et al., High-Level Expression of AmpC β-Lactamase Due to Insertion of Nucleotides between −10 and −35 Promoter Sequences in Escherichia coli Clinical Isolates: Cases Not Responsive to Extended-Spectrum-Cephalosporin Treatment. Antimicrobial Agents and Chemotherapy, 2003. 47(7): p. 2138–2144.

32. Tracz, D.M., et al., Increase in ampC promoter strength due to mutations and deletion of the attenuator in a clinical isolate of cefoxitin-resistant Escherichia coli as determined by RT-PCR. J Antimicrob Chemother, 2005. 55(5): p. 768–72.

33. Tracz, D.M., et al., ampC gene expression in promoter mutants of cefoxitin-resistant Escherichia coli clinical isolates. FEMS Microbiol Lett, 2007. 270(2): p. 265–71.

34. Caroff, N., et al., Analysis of the Effects of −42 and −32 ampC Promoter Mutations in Clinical Isolates of Escherichia coli Hyperproducing AmpC. Journal of Antimicrobial Chemotherapy, 2000. 45: p. 783–788.

35. Jaurin, B., T. Grundstrom, and S. Normark, Sequence elements determining ampC promoter strength in E. coli. The EMBO Journal 1982. 1(7): p. 875–881.

36. Srinivasan, V.B., et al., Role of the two component signal transduction system CpxAR in conferring cefepime and chloramphenicol resistance in Klebsiella pneumoniae NTUH-K2044. PLoS One, 2012. 7(4): p. e33777.

37. Batchelor, E., et al., The Escherichia coli CpxA-CpxR envelope stress response system regulates expression of the porins ompF and ompC. J Bacteriol, 2005. 187(16): p. 5723–31.

38. Cianfanelli, F.R., et al., VgrG and PAAR Proteins Define Distinct Versions of a Functional Type VI Secretion System. PLoS Pathog, 2016. 12(6): p. e1005735.

39. Nicoloff, H., et al., The high prevalence of antibiotic heteroresistance in pathogenic bacteria is mainly caused by gene amplification. Nat Microbiol, 2019. 4(3): p. 504–514.

40. El Meouche, I., Y. Siu, and M.J. Dunlop, Stochastic expression of a multiple antibiotic resistance activator confers transient resistance in single cells. Sci Rep, 2016. 6: p. p.

41. Foxman, B., The epidemiology of urinary tract infection. Nat Rev Urol, 2010. 7(12): p. 653–60.

42. Anderson, G.G., et al., Intracellular Bacterial Biofilm-Like Pods in Urinary Tract Infection. Science, 2003. 301(5629): p. 105–107.

43. Podnecky, N.L., et al., Conserved collateral antibiotic susceptibility networks in diverse clinical strains of Escherichia coli. Nat Commun, 2018. 9(1): p. 3673.

44. Suzuki, S., T. Horinouchi, and C. Furusawa, Prediction of antibiotic resistance by gene expression profiles. Nat Commun, 2014. 5: p. 5792.

45. Dean, C.R., et al., Mode of Action of the Monobactam LYS228 and Mechanisms Decreasing In Vitro Susceptibility in Escherichia coli and Klebsiella pneumoniae. Antimicrob Agents Chemother, 2018. 62(10): p. 01200–18.

46. Wang, J., et al., The role of the type VI secretion system vgrG gene in the virulence and antimicrobial resistance of Acinetobacter baumannii ATCC 19606. PLoS One, 2018. 13(2): p. e0192288.

47. The International Organization for Standardization. ISO 20776-1:2006 Clinical laboratory testing and in vitro diagnostic test systems — Susceptibility testing of infectious agents and evaluation of performance of antimicrobial susceptibility test devices. Part 1: Reference method for testing the in vitro activity of antimicrobial agents against rapidly growing aerobic bacteria involved in infectious diseases. 2006 The International Organization for Standardization. https://www.iso.org/obp/ui/#iso:std:iso:20776:-1:ed-1:v1:en.

48. Hu, Y., et al., A new approach for the discovery of antibiotics by targeting non-multiplying bacteria: a novel topical antibiotic for staphylococcal infections. PLoS One, 2010. 5(7): p. e11818.

49. Birosova, L. and M. Mikulasova, Development of triclosan and antibiotic resistance in Salmonella enterica serovar Typhimurium. J Med Microbiol, 2009. 58(Pt 4): p. 436–41.

50. Kavanagh, A., et al., Effects of Microplate Type and Broth Additives on Microdilution MIC Susceptibility Assays. Antimicrobial Agents and Chemotherapy, 2018. 63(1): p. e01760–18.

51. Sabir, N., et al., Bacterial biofilm-based catheter-associated urinary tract infections: Causative pathogens and antibiotic resistance. Am J Infect Control, 2017. 45(10): p. 1101–1105.

52. Manges, A.R., et al., Widespread distribution of urinary tract infections caused by a multidrug-resistant Escherichia coli clonal group. The New England Journal of Medicine, 2001. 345(14): p. 1007–1013.

53. Yamaji, R., et al., Persistent Pandemic Lineages of Uropathogenic Escherichia coli in a College Community from 1999) to 2017. Journal of Clinical Microbiology, 2018. 56(4): p. e01834–17.

